# Mutagenesis of human cytomegalovirus glycoprotein L disproportionately disrupts gH/gL/gO over gH/gL/pUL128-131

**DOI:** 10.1101/2021.04.09.439256

**Authors:** Eric P. Schultz, Qin Yu, Cora Stegmann, Le Zhang Day, Jean-Marc Lanchy, Brent J. Ryckman

**Author notes:** Corresponding author: Dr. Eric P. Schultz, Division of Biological Sciences, Interdisciplinary Sciences Building Rm. 221, University of Montana, Missoula, MT 59812.

## Abstract

Cell-free and cell-to-cell spread of herpesviruses involves a core fusion apparatus comprised of the fusion protein glycoprotein B (gB) and the regulatory factor gH/gL. The human cytomegalovirus (HCMV) gH/gL/gO and gH/gL/pUL128-131 facilitate spread in different cell types. The gO and pUL128-131 components bind distinct receptors, but the how the gH/gL portion of the complexes functionally compare is not understood. We previously characterized a panel of gL mutants by transient expression and showed that many were impaired for gH/gL-gB dependent cell-cell fusion, but were still able to form gH/gL/pUL128-131 and induce receptor-interference. Here, the gL mutants were engineered into the HCMV BAC clones TB40/e-BAC4 (TB), TR and Merlin (ME), which differ in their utilization of the two complexes for entry and spread. Several of the gL mutations disproportionately impacted gH/gL/gO-dependent entry and spread over gH/gL/pUL128-131 processes. Effects of some mutants could be explained by impaired gH/gL/gO assembly, but other mutants impacted gH/gL/gO function. Soluble gH/gL/gO containing the L201 mutant failed to block HCMV infection despite unimpaired binding to PDGFRα, indicating the existence of other important gH/gL/gO receptors. Another mutant (L139) enhanced the gH/gL/gO-dependent cell-free spread of TR, suggesting a “hyperactive” gH/gL/gO. Recently published crystallography and cryo-EM studies suggest structural conservation of the gH/gL underlying gH/gL/gO and gH/gL/pUL128-131. However, our data suggest important differences in the gH/gL of the two complexes and support a model in which gH/gL/gO can provide an activation signal for gB.

**IMPORTANCE:** The endemic *beta*-herpesvirus HCMV circulates in human populations as a complex mixture of genetically distinct variants, establishes lifelong persistent infections, and causes significant disease in neonates and immunocompromised adults. This study capitalizes on our recent characterizations of three genetically distinct HCMV BAC clones to discern the functions of the envelope glycoprotein complexes gH/gL/gO and gH/gL/pUL128-13, which are promising vaccine targets that share the herpesvirus core fusion apparatus component, gH/gL. Mutations in the shared gL subunit disproportionally affected gH/gL/gO, demonstrating mechanistic differences between the two complexes and may provide a basis for more refined evaluations of neutralizing antibodies.

## INTRODUCTION

Next-generation sequencing and genomics studies have presented a complex and dynamic picture of human cytomegalovirus (HCMV) genetics in human populations (1–9). Twenty-one of HCMV’s 165 canonical genes show relatively higher levels of sequence diversity, existing as 2-14 “genotypes”, or “alleles” distributed throughout the remainder of a more highly conserved genome. Genetic signatures of recombination suggest that these variable alleles can be shuffled into a very large number of individual haplotypes and individuals can harbor complex, dynamic mixtures of haplotypes. Given these observations, it is not surprising that clinical specimens can contain complex mixtures of HCMV haplotypes. The observed adaptations to laboratory culture conditions may involve selection or random sampling of preexisting haplotypes in addition to the arising of *de novo* mutations (10–14). While the modern practices of capturing HCMV haplotypes as bacterial artificial chromosome (BAC) clones provide stability and convenient genetic manipulation approaches, it also may obscure the significance of complex genetic diversity to the mechanisms of virus replication.

Like other herpesviruses, HCMV uses a core fusion apparatus comprised of the fusion protein glycoprotein B (gB) and the regulatory factor gH/gL to spread via entry of extracellular viruses (“cell-free”), or via direct cell-to-cell spread (reviewed in (15)). HCMV gH/gL is bound by either gO, or the UL128-131 proteins to form complexes that influence cell-type tropism through a variety of potential receptor interactions. Efficient entry into all cell types within the broad tropism range of HCMV depends on gH/gL/gO, which has been shown to bind platelet-derived growth factor receptor alpha (PDGFRα) and transforming growth factor beta receptor type 3 (TGFβRIII) (16–20). Entry into epithelial and endothelial cells, and some leukocytes is greatly enhanced by gH/gL/pUL128-131, which can bind to neurophilin-2 (NRP-2), olfactory receptor (OR) 14I1, and β1 and β3 integrins, but gH/gL/pUL128-131 is dispensable for entry into fibroblasts and neuronal cells (21–28). Either gH/gL complex can suffice for direct cell-to-cell spread in fibroblasts cultures, whereas gH/gL/pUL128-131 is required in epithelial and endothelial cell cultures (16, 29–31).

We and others have characterized how the HCMV BAC clones TB40/e (TB), TR, and Merlin (ME) differ in their expression of gH/gL/gO and gH/gL/pUL128-131 and how this influences entry and spread in fibroblasts and epithelial cells. Extracellular TB and TR virions contain far more gH/gL/gO than gH/gL/pUL128-131, whereas ME has overall less total gH/gL and this is mostly in the form of gH/gL/pUL128-131 (19, 32). The ME BAC clone used in our studies was engineered by Stanton et al. with tetracycline (Tet) operator sequences in the UL131 promoter such that replication in cells expressing the Tet repressor protein (TetR) yields virions with greatly diminished gH/gL/pUL128-131 and slightly more gH/gL/gO (19, 33). Specific infectivity of this set of BAC clones for fibroblasts and epithelial cells does not strictly correlate with the abundances of the gH/gL complexes, indicating important contributions from other variable viral factors (19). Likewise, direct cell-to-cell spread efficiency is not solely determined by gH/gL complexes (34). In fibroblasts, TB is highly efficient for cell free spread and particularly poor at cell-to-cell spread, ME is highly efficient at cell-to-cell, but not cell free spread, and TR utilizes both spread modes more evenly, but less efficiently. The efficiency of cell-to-cell spread by ME in fibroblasts was not impaired by Tet-repression of gH/gL/pUL128-131, indicating the contribution of mechanisms beyond those provided by gH/gL complexes. In epithelial cells, ME is far more efficient at spread than either TB or TR, and this was impaired by Tet-repression of gH/gL/pUL128-131 (31, 34). However, observations that the specific paring of the variable alleles of gH and gO can impact the efficiency of spread in epithelial cells suggests that gH/gL/gO can also contribute (35, 36). The RL13 protein has been suggested to selectively restrict cell-free spread in favor of cell-to-cell spread (10, 33, 37). However, our analyses suggest that pRL13 tempers spread by either mode (34). Finally, the mechanism of spread may also be influenced by the nature of the producer cell type itself. Producer cell effects on the expression of gH/gL complexes have been described, but not analyzed in detail (34, 38).

The basic models of herpesvirus membrane fusion suggest that receptor binding by gH/gL or by accessory proteins like gD of herpes simplex virus (HSV) or gp42 or Epstein-Barr virus (EBV) expose surfaces on gH/gL that can interaction with gB and promoting fusion (reviewed in (15)). It is not yet clear whether such a model applies also to HCMV gH/gL/gO, gH/gL/pUL128-131, or the more recently described gH/pUL116 complex (39). Transient expression of just gH/gL and gB is sufficient to drive cell-cell fusion but this may not recapitulate the regulation of fusion during viral entry (40). In a previous report, we used replication-defective adenovirus expression vectors to characterize a library of charged cluster-to-alanine (CCTA) mutants of HCMV gH and gL with the hypothesis that some might mechanistically distinguish gH/gL/gO and gH/gL/pUL128-131 (41). None of the mutations disrupted the formation of gH/gL dimers, but most of these were impaired in the gH/gL-gB cell-cell fusion assay. Most could still support the assembly of gH/gL/pUL128-131 capable of inducing receptor-interference in epithelial cells, but assembly and function of gH/gL/gO was not addressed. In the current report, we exploited the well characterized, and highly specialized spread properties of the BAC clones TB, TR, and ME to evaluate the effects of the gL mutations on the functions of gH/gL/gO and gH/gL/pUL128-131. Data are presented indicating that several of the gL mutants disproportionally impair the assembly and function of gH/gL/gO over gH/gL/pUL128-131, and implications of these results for the mechanisms by which these complexes facilitate entry and spread are discussed.

## RESULTS

### Effects of HCMV gL CCTA mutations on assembly of soluble gH/gL/gO and gH/gL/pUL128-131

Figure 1A lists all amino acids mutated in gL with the numerical designations referring to the first amino acid of each cluster. The recently reported cryo-EM derived structure of the gH/gL/gO trimer complex shows that gL forms a bridge between gH and gO (Fig 1B) (42). Mapping the CCTA mutations onto this model predicts L46 and L63 to be solvent exposed, L139 and L156 making direct interactions with gO, L201 interfacing the gH- and gO-binding regions, and L244 and L256 involved in core interactions with gH (Fig 1C). To evaluate the effect of these gL mutations on assembly of gH/gL/gO, adenovirus (Ad) expression vectors were used. When expressed alone, all gL mutants accumulated to detectable steady-state levels within cell extracts (Fig 2A). The small differences in band intensities observed might indicate differences in stability or turnover rates, but also could reflect differences in immunoblot transfer efficiency or antibody reactivity due to the various mutations of charged amino acids. When cells were transduced with Ad vectors encoding soluble gH (sgH) (43) plus the indicated WT or mutant gL, and either gO or pUL128-131, all culture supernatants contained comparable amounts of sgH/gL/gO complexes except for those expressing L156, which displayed significantly less sgH/gL/gO, but similar levels of sgH (Fig 2B). Because no gH was secreted in the absence of gL, the gH present in the L156 lane likely represented sgH/gL-sgH/gL homodimers, as previously described (44). When sgH and WT or mutant gLs were expressed with pUL128, pUL130 and pUL131, all supernatants contained comparable amounts of disulfide-linked sgH/gL/pUL128 complexes (Fig 2C top panel), consistent with our previous study (41). Note that the pUL130 and pUL131 proteins are disulfide-linked to each other, but not covalently bound to gH/gL/pUL128, and thus dissociate from the complex during SDS-PAGE (44–46). To confirm assembly of complete gH/gL/pUL128-131, supernatants were also analyzed by immunoblot for pUL130/131. Complexes formed with mutants L46, L63, L139, and L201 contained comparable amounts of pUL130/131 as those formed with WT gL. However, less pUL130/131 was detected with L156, L244 and L256. Given that Ryckman et al. showed that secretion of soluble gH/gL/pUL128, lacking pUL130/131 was highly inefficient (45), it seems likely that L156, L244 and L256 facilitate assembly and secretion of intact gH/gL/pUL128-131 complexes, but that after secretion, the pUL130/131 are more prone to dissociation, explaining the normal levels of gH/gL/pUL128 but the reduced pUL130/131 detected.

**Figure 1.**
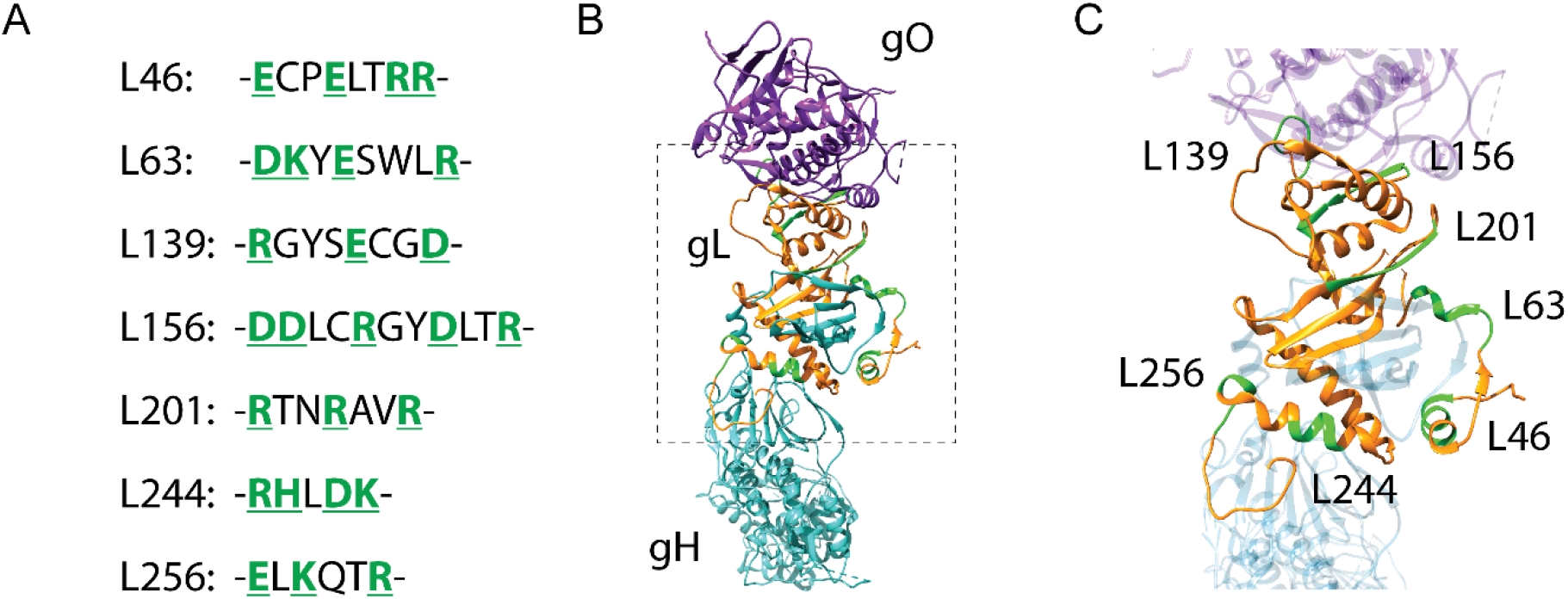
Locations and specific AA changes for the gL CCTA mutations. (A) Specific amino acid residues changed for gL mutants. Mutants are designated by the starting residue of the cluster and the specific residues changed to alanine are indicated (green). (B) 3-D representation of the gH/gL/gO complex of HCMV (7LBE, Kschonsak et al.). Glycoprotein L (gL, orange) interacts with both gH (cyan) and gO (purple) through separate binding domains. (C) Closer look at the locations of specific CCTA mutations in gL (green) and their proximity to gH and gO.

**Figure 2.**
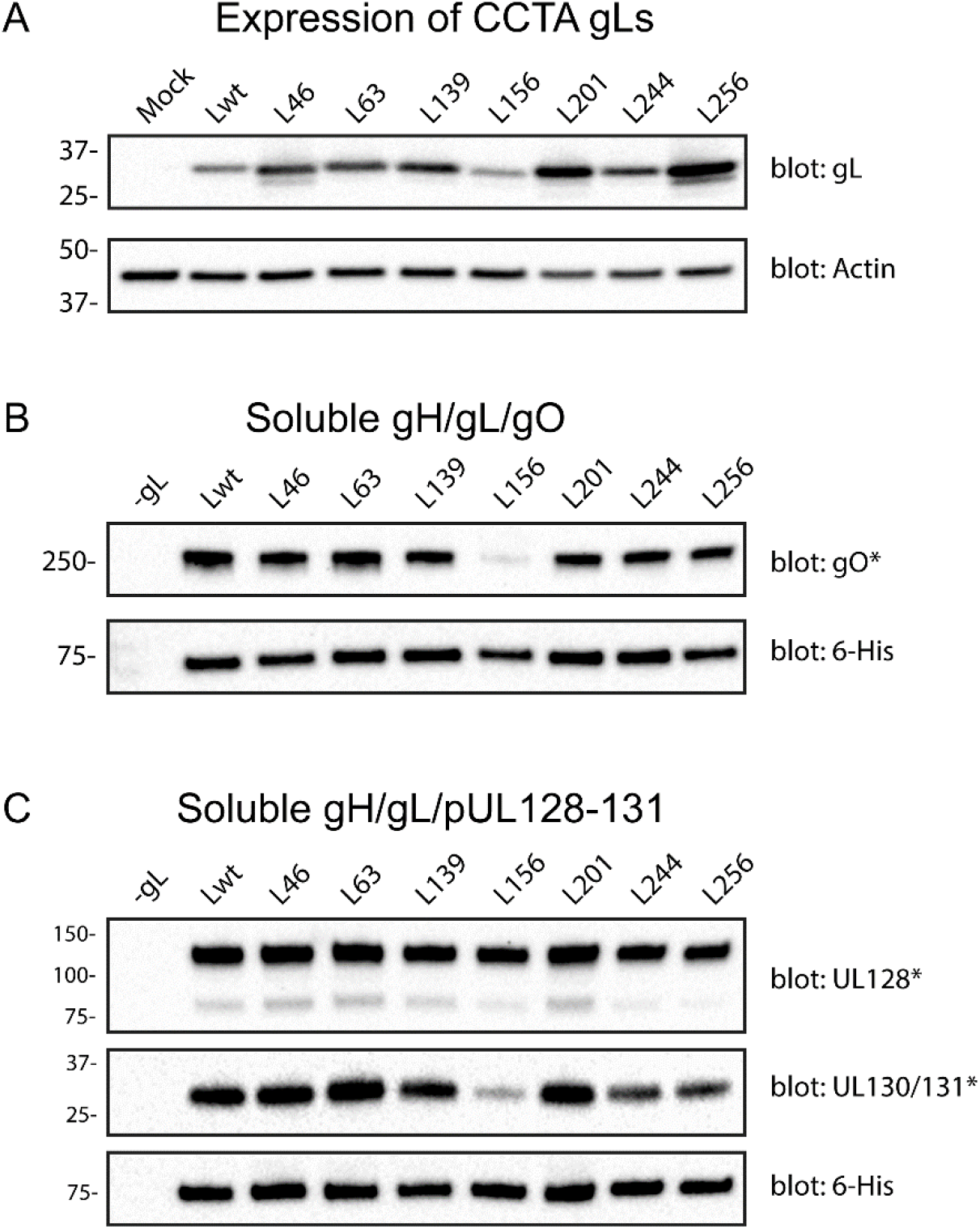
Expression of soluble gH/gL complexes containing CCTA gL mutations. (A) Cells were infected with Ad vectors expressing wild type and mutant gLs. Cell extracts were analyzed by SDS-PAGE followed by immunoblot for gL and actin. (B-C) Cells were infected with Ad vectors expressing soluble gH-6His, gL (wild type or indicated mutants), and gO (B) or pUL128, pUL130, and pUL131 (C). Soluble gH/gL complexes were enriched using Ni-NTA agarose resin and analyzed by SDS-PAGE followed by immunoblot for gH (6His), gO, UL128, or UL130/UL131. SDS-PAGE separations indicated with an asterisk were performed under non-reducing (-DTT) conditions to preserve disulfide linkages of gH/gL complexes.

### Soluble gH/gL/gO complexes containing mutant gL bind the receptor PDGFRα, but L201 fails to block HCMV infection

Others have shown that sgH/gL/gO complexes can block HCMV infection, and this is generally attributed to the saturation of PDGFRα on the cell surface (20, 47, 48). Soluble gH/gL/gO complexes containing mutants L46, L63, L139, L244, and L256 were able to block HCMV infection with similar potency as wild type however complexes containing mutations L156 and L201 were ineffective (Fig 3A). This was not surprising for L156, since this mutation caused a dramatic reduction in the assembly of gH/gL/gO (Fig 2B). In contrast, the reduced HCMV blocking by sgH/gL201/gO could not be explained by effects on complex formation or stability, so we tested the mutant gH/gL/gO complexes for direct interaction with PDGFRα-Fc by ELISA. Surprisingly, all mutant gH/gL/gO complexes, including L201 bound comparably to PDGFRα-Fc (except L156, which as noted above, failed to produce intact gH/gL/gO complexes) (Fig 3B). ELISA results were corroborated by an affinity pull-down approach where soluble gH/gL/gO complexes and PDGFRα-Fc were incubated together, then captured by Ni-NTA enrichment of the sol gH-6His tag and analyzed by immunoblot (Fig 3C). EC_50_ values for both HCMV inhibition and PDGFRα-Fc binding are presented in Table 1. The soluble L201 trimer resulted in similar EC_50_ values for both inhibition and PDGFRα-Fc binding despite a substantial reduction in maximal HCMV inhibition (31% compared to 81% for WT). This indicates that the inability of sgH/gL201/gO complexes to block HCMV infection is not due to a lack of PDGFRα binding. Thus, while engagement of PDGFRα by gH/gL/gO is required for efficient entry of HCMV, gH/gL/gO likely interacts with other critical cell-surface proteins either up- or downstream of PDGFRα engagement, and this may be the basis for the observed blocking of infection by sgH/gL/gO.

**Table 1:** Analysis of soluble gH/gL/gO mutants for HCMV inhibition and PDGFRα-Fc binding

| gH/gL/gO<br>(gL mutant) | HCMV Inhibition<br>(EC<sub>50</sub>, dilution) | p-value<sup>a</sup> | Max Inhibition<br>(%) | p-value<sup>a</sup> | PDGFRα-Fc Binding<br>(EC<sub>50</sub>, ng/mL) | p-value<sup>a</sup> |
| --- | --- | --- | --- | --- | --- | --- |
| Blank | n/a | n/a | none | n/a | 3702 ± 997.2 | <0.0001 |
| HL | n/a | n/a | none | n/a | 1285 ± 100.9 | <0.0001 |
| WT | 0.019 ± 0.001 | n/a | 80.8 ± 3.6 | n/a | 125.7 ± 16.88 | n/a |
| L46 | 0.029 ± 0.001 | 0.0145 | 82.1 ± 1.1 | 0.9662 | 180.4 ± 10.78 | 0.9222 |
| L63 | 0.016 ± 0.002 | 0.5751 | 80.7 ± 1.6 | 0.9998 | 175.0 ± 13.42 | 0.9529 |
| L139 | 0.016 ± 0.003 | 0.5607 | 78.7 ± 2.7 | 0.7930 | 168.0 ± 4.822 | 0.9795 |
| L156 | n/a | n/a | none | n/a | 1298 ± 66.12 | <0.0001 |
| L201 | 0.021 ± 0.001 | 0.9743 | 31.0 ± 1.5 | <0.0001 | 219.1 ± 25.69 | 0.5200 |
| L244 | 0.015 ± 0.001 | 0.3664 | 75.3 ± 1.1 | 0.0702 | 190.1 ± 18.31 | 0.8444 |
| L256 | 0.014 ± 0.003 | 0.1556 | 65.3 ± 3.9 | <0.0001 | 185.5 ± 13.62 | 0.8843 |

<sup>a</sup> ANOVA, Dunnett's multiple comparisons to WT

**Figure 3.**
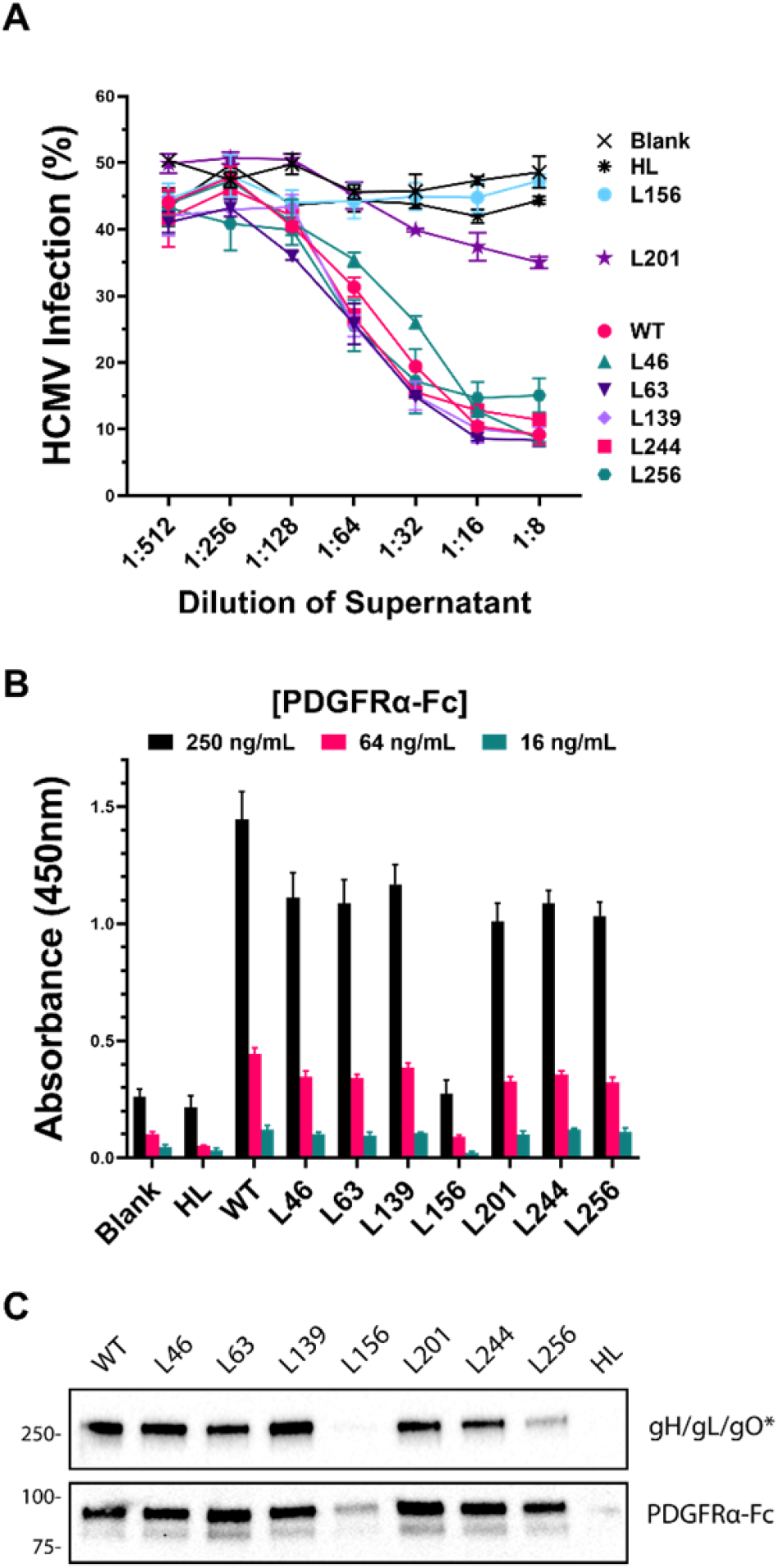
Effect of gL mutations on HCMV inhibition and PDGFRα binding. (A) Fibroblasts were incubated at 4°C with dilutions of supernatants from cells expressing the indicated combinations of gH-6His, gL (or mutant gL), and gO, then inoculated with HCMV at 37°C and infection was measured 2 d.p.i. by flow cytometry. (B) Ni-NTA ELISA plates were coated with soluble gH/gL/gO mutants and incubated with increasing concentrations of PDGFRα-Fc. Binding was detected following incubation with an HRP-conjugated anti-human antibody and measured by colorimetric analysis. (C) Soluble gH/gL/gO mutants were incubated at 37°C with PDGFRα-Fc and then complexes were pulled down with Ni-NTA agarose and analyzed by SDS-PAGE followed by immunoblot.

### Effects of CCTA gL mutations on spread efficiency and infectivity of HCMV

In a previous report we demonstrated that the commonly studied HCMV BAC clones Merlin (ME) and TB40/e-BAC4 (TB) spread in fibroblasts cultures with very similar efficiencies over 12 days (34). However, whereas ME is highly specialized for the cell-to-cell mode and produces tightly localized foci, TB is highly specialized for the cell-free mode and produces more diffuse foci. In contrast, the TR BAC clone spreads less efficiently than either TB or ME, but utilizes both modes more evenly. Thus, TR was chosen as the genetic background for the initial characterizations of the gL mutants. A gL-complementing cell line was used to mitigate potential reversions and second-site suppressor mutations during mutant virus propagation. Constitutive expression of gL in these cells was lower than in HCMV-infected cells, but was enhanced by HCMV infection, demonstrating that the gL-expressing cells remain susceptible to infection (Fig 4A). Despite the lower expression of gL compared to WT infection, the gL-nHDF cells efficiently complemented the severe spread defect of TR_UL115stop (TRΔgL) (Fig 4B). Using viruses grown in complementing gL-nHDFs, we found that all gL mutants were expressed at lower steady-state levels compared to wild type gL during HCMV infection of non-complementing cells (Fig 4C). This result was different than the analysis of Ad vector-expressed sgH/gL complexes (Fig 2), suggesting that these gL mutations influence the mechanism of gH/gL complex assembly in HCMV infected cells, which involves other viral proteins such as pUL148, pUL116, and pUS16 (43, 49, 50).

**Figure 4.**
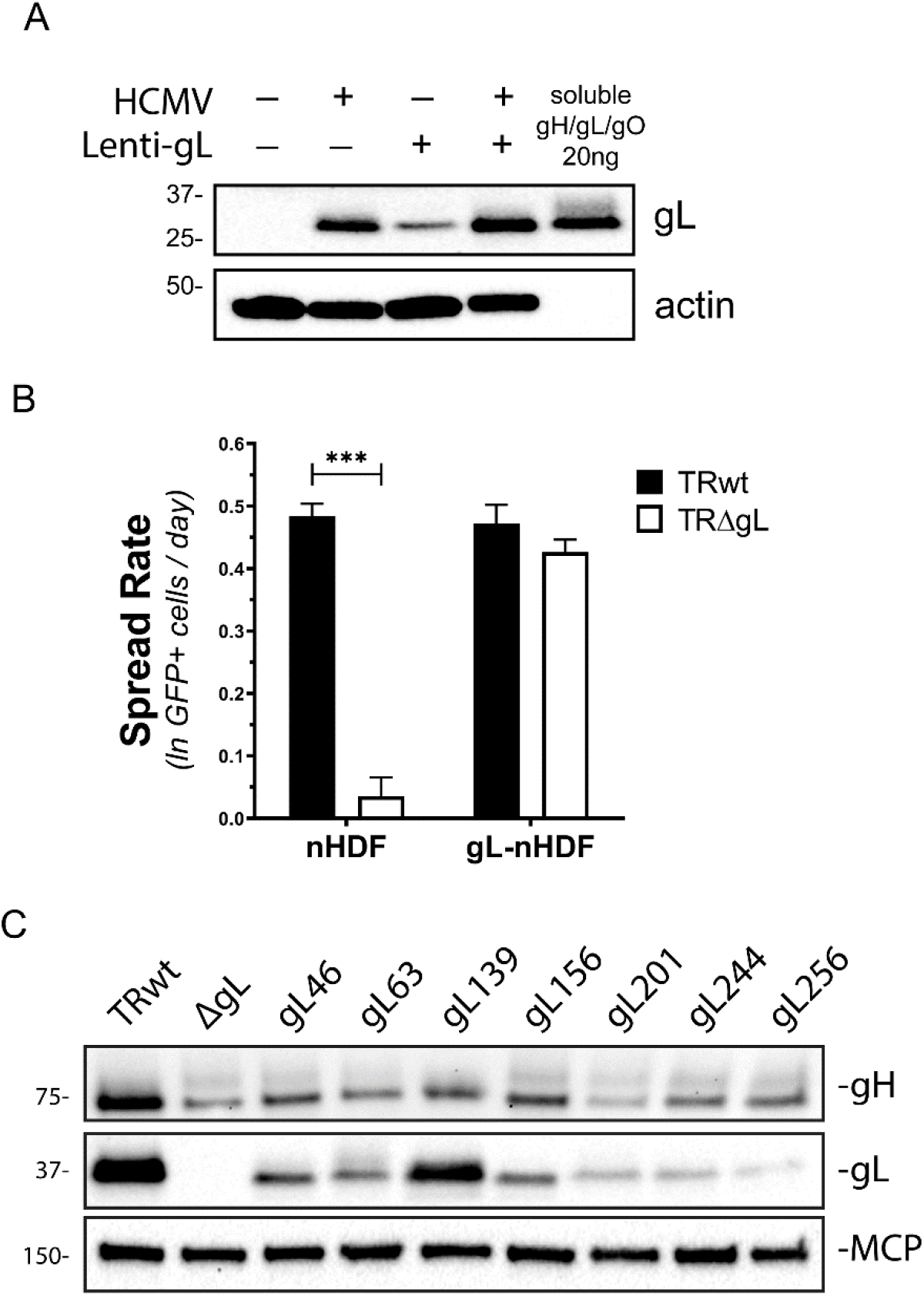
Evaluation of gL-expressing fibroblasts for complementation of CCTA gL mutant HCMV. (A) Primary nHDFs or those transduced with lentiviral vectors encoding UL115 were infected with HCMV (MOI 1) or mock infected and cells were extracted at 5 d.p.i. Samples were analyzed by SDS-PAGE followed by immunoblot for gL and actin. 20ng of purified gH/gL/gO was loaded for comparison. (B) Normal and gL-transduced nHDF cells were infected with either TRwt or TRΔgL HCMV and spread was monitored over 12 days by flow cytometry. Results shown are average spread rates of three experiments and error bars represent standard deviation. P-values calculated using paired t-tests (***<0.001). (C) nHDF cells infected with complemented gL mutants were analyzed by SDS-PAGE followed by immunoblot for gH, gL, and MCP.

Non-complementing nHDFs were infected with complemented TR-based gL mutants and foci were evaluated 12 days post infection using fluorescence microscopy (Fig 5A). As expected, TRΔgL failed to spread beyond the initial infected cells. However, the gL mutants spread to form foci of varying sizes and patterns. L139 mutants generated more diffuse foci than the parental TR while L201, L244, and L256 foci were notably smaller and more compact. L156 mutants generated very small foci, typically consisting of only 2-3 cells but were distinctly larger than ΔgL. For a more rigorous analysis, spread rates were determined using a previously described quantitative flow cytometry approach (Fig 5B)(34). Spread rates for the mutants closely corresponded with their respective focal appearance, with L46 and L63 being like wild type and L201, L244, and L256 spreading at a slower rate. L139 spread significantly faster than wild type while L156 spread only marginally better than the ΔgL mutant.

**Figure 5.**
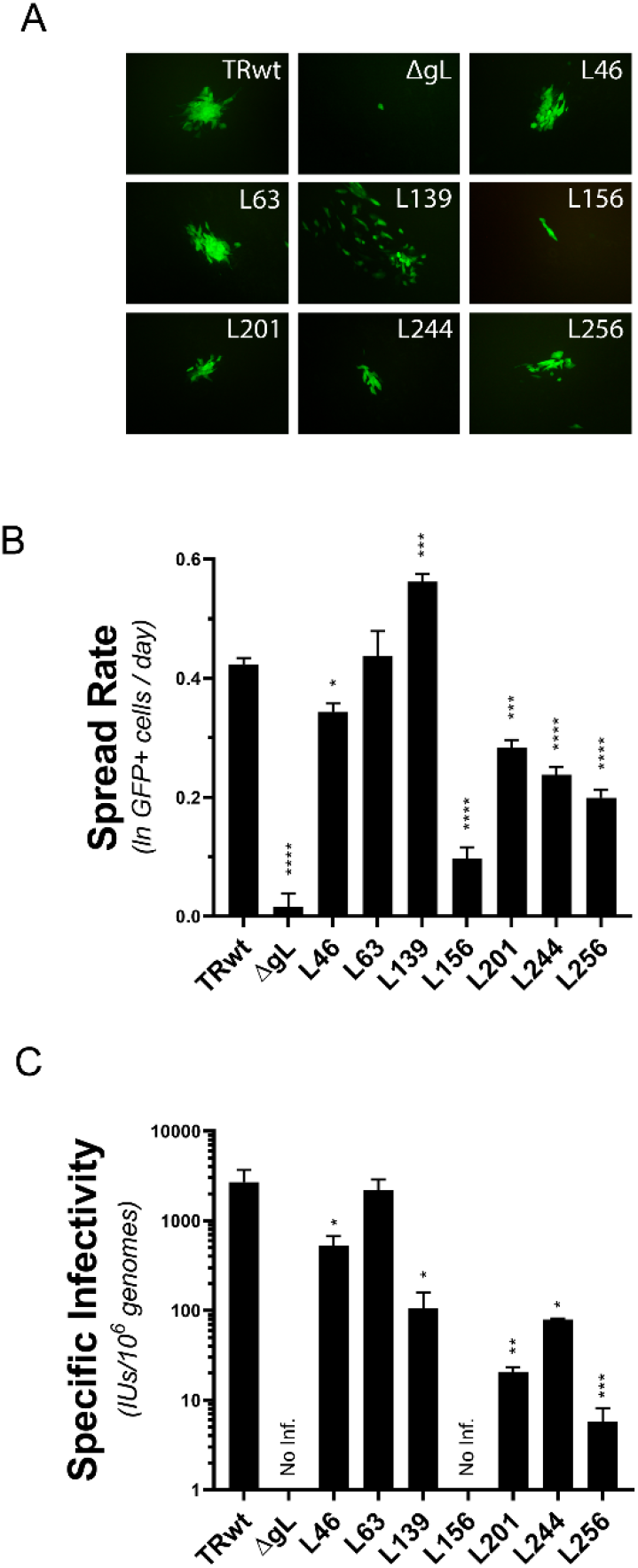
CCTA gL mutations affect the spread and cell-free infectivity of HCMV strain TR. (A) Normal nHDF cells were inoculated with complemented gL mutant HCMVs and foci were analyzed 12 d.p.i. using fluorescence microscopy. (B) Spread efficiency of the gL mutants was monitored over 12 days using flow cytometry. Average rates for three experiments are shown. (C) Normal nHDFs were infected with complemented gL mutant HCMVs and supernatants were harvested 8 d.p.i. HCMV genomes/mL was measured by qPCR and IUs were determined on nHDF cells using flow cytometry. Average specific infectivities of three preparations for each mutant are shown. Viruses for which no infectivity could be measured are labelled “No Inf”. (B-C) Error bars represent standard deviation and P values were calculated using ANOVA with Dunnett’s multiple comparison test to WT (*<0.05, **<0.01, ***<0.001, ****<0.0001).

The observed diffuse focal pattern and increased spread rate of TR_L139 suggested an enhanced cell-free mechanism of spread, which generally correlates with specific infectivity (IUs/genome) of the cell-free virions (34). Specific infectivity of TR_ L63 was comparable to the parental TR, while TR_L46, L139, L201, L244, and L256 were each moderately impaired, and TR_L156 was severely impaired (Fig 5C). Thus, cell-free infectivity and spread efficiency correlated for most mutants, but for L139, the increased spread rate and diffuse focal pattern was despite a reduced cell-free infectivity. An explanation for this miscorrelation may be that the specific infectivity assays involved harvesting and storage of supernatant virus, which is likely more demanding on the structural integrity of the virions compared to the spread assays where the newly produced virions have more immediate access to new host cells. Since sgH/gL139/gO complexes were not grossly unstable (Fig. 2B), it may be that the L139 mutation renders the active conformation of gH/gL/gO more labile, and this leads to more loss of infectivity during harvesting and storage of the virus prior to specific infectivity analyses.

### Effects of CCTA gL mutants on spread by extracellular virus

To more specifically address the cell-free mode of spread, mutants ΔgL, L46, L139, L156, and L201 were engineered into the TB BAC clone, which is highly specialized to the cell-free mode of spread (34). While TB_L46 and L139 spread at rates indistinguishable from TBwt, TB_L156 was highly impaired, and L201 was moderately impaired (Fig 6A). Given the reliance of TB on highly infectious extracellular virus for efficient spread, these results were largely explained by the findings that TB_L46 and L139 virions were as infectious as TBwt, whereas TB_L156 and TL201 virions were noninfectious (Fig 6B). To assess whether these infectivity characteristics could be explained by the amounts of gH/gL complexes in the TB virions, non-reducing immunoblots were performed as before (19). Consistent with previous analyses, extracts of TBwt virions contained gH/gL predominately in the form of gH/gL/gO, and contained very little gH/gL/pUL128-131 (Fig. 6C). TB_L139 had similar levels of gH/gL complexes as the parental TBwt, whereas TB_L46 was reduced in gH/gL/gO, and TB_L201 reduced even further. Neither TB_L46 nor L201 had an offsetting increase in gH/gL/pUL128-131. TB_L156 extracts contained a gL species that migrated markedly faster than the gH/gL/gO bands of the other viruses. Stripping the gL antibodies from the blots and re-probing with anti-gO antibodies demonstrated that this gL species was not gH/gL/gO, and this was consistent with the inability of L156 to support assembly of sgH/gL/gO during Ad vector expression (Fig 2B). The nature of the faster migrating gL species in TB_L156 remains unclear but possibilities include; 1) gH/gL-gH/gL “dimer of dimers”, which can form through Cys144 of gL that would normally bind to either pUL128 or gO (44, 51), or 2) gH/gL bound by other cellular or viral “chaperone-like” proteins.

**Figure 6.**
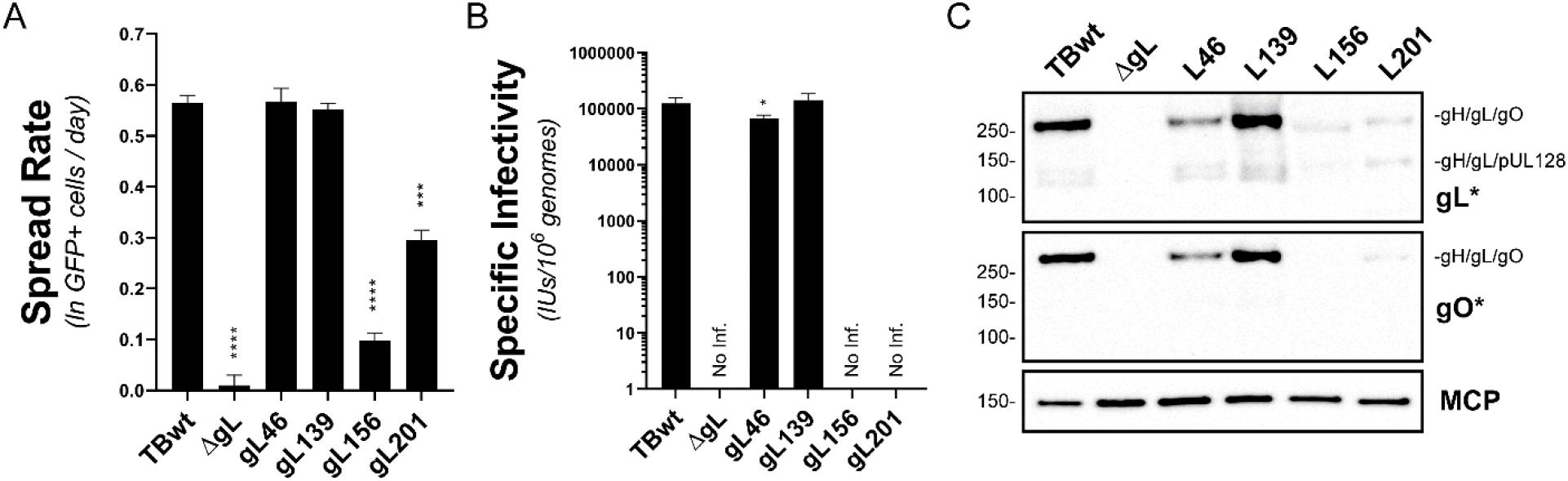
Effect of gL mutagenesis on spread of HCMV strain TB. (A) nHDF cells were infected at MOI 0.001 with complemented TB gL mutants and spread efficiency was monitored over 12 days by flow cytometry. (B) nHDF cells were infected at MOI 1 with indicated TB gL mutants and virus-containing supernatants were collected 7 d.p.i. HCMV genomes/mL was measured by qPCR and IUs were determined on nHDF cells using flow cytometry. Average specific infectivities of three preparations for each mutant are shown. Viruses for which no infectivity could be measured are labelled “No Inf”. (C) nHDF cells were infected with HCMV strain TB (WT or indicated gL mutants) and virus-containing supernatants were collected 7 d.p.i. Virions were analyzed by SDS-PAGE followed by immunoblot for gL, gO, and MCP. Asterisks indicate SDS-PAGE performed under non-reducing (-DTT) conditions. (A-B) All error bars represent standard deviation and P values were calculated using ANOVA with Dunnett’s multiple comparison test to WT (*<0.05, **<0.01, ***<0.001, ****<0.0001).

The lack of gH/gL/gO in TB_L156 virions provides a compelling explanation for the lack of observed infectivity. By contrast, infectivity of TB_201 virions was also undetectable, but this was not as easily attributed to the reduced gH/gL/gO in the virion for a number of reasons. First, whereas TB_156 virions was devoid of gH/gL/gO, TB_L201 virions clearly contain gH/gL/gO. This difference is likely the basis of why TB_L201 spread at 3 times the rate of TB_L156 (Fig 6A). Second, TB_L46 was also reduced in gH/gL/gO but was nearly identical to the parental TB in both spread and infectivity (Fig. 6A and B). The dramatic differences in infectivity and spread between TB_L46 and TB_L201 seem out of proportion with the difference in gH/gL/gO abundance, suggesting that the L201 mutation impairs not only gH/gL/gO assembly, but also function. In support of this interpretation, L201 was the only gL mutant to yield sgH/gL/gO that failed to block HCMV infection (Fig 3A).

The severe impact of L201 on the infectivity of TB and the lack of impaired infectivity of TB_L139 stand in contrast to the observations of these mutations in the TR background. Non-mutually exclusive explanations for these discrepancies include; 1) impacts of these gL mutations on the gH/gL/gO complexes may be dependent on genetic differences in the gH and gO encoded by these strains and, 2) the relative contribution of gH/gL/gO to the observed infectivity of these strains may be different due to functional variations associated with other entry glycoproteins such as gB or gM/gN, or even other early infection processes such as nuclear translocation or gene expression.

### Effects of CCTA gL mutants on direct cell-to-cell spread

The effects of gL mutations on cell-to-cell spread were assessed using the cell-to-cell specialist BAC clone, ME (34). Mutants L46 and L139 had little or no effect of spread of ME in fibroblasts, whereas L156 and L201 reduced spread by 2- and 1.5-fold, respectively (Fig 7A). Previous studies of gO-null ME suggested that cell-to-cell spread of ME could be facilitated by its robust expression of gH/gL/pUL128-131, independent of gH/gL/gO (29). However, given the effects of L156 and L201 on gH/gL/gO indicted above, the impaired spread of ME_L156 and L201 suggested a contribution of gH/gL/gO to ME spread, or effects on the gH/gL/pUL128-131. Alternative cell culture systems were used to distinguish the contribution of the two gH/gL complexes. The ME BAC clone used for these studies was engineered with tetracycline-operator sequences in the UL131 transcriptional promoter (33). In fibroblasts expressing the tetracycline repressor protein (TetR), the assembly of gH/gL/pUL128-131 is repressed and spread of ME is dependent on gH/gL/gO (19, 34). In these TetR expressing cells, spread of ME_L46, L156, and L201 mutations was more impaired compared to regular fibroblasts (Fig 7B). Conversely, none of the gL mutations had much effect on spread of ME in epithelial cells, where spread is highly dependent on gH/gL/pUL128-131 (Fig 7C). Together these results suggests that these gL mutations disproportionately affect the function of gH/gL/gO over gH/gL/pUL128-131 in cell-to-cell spread.

**Figure 7.**
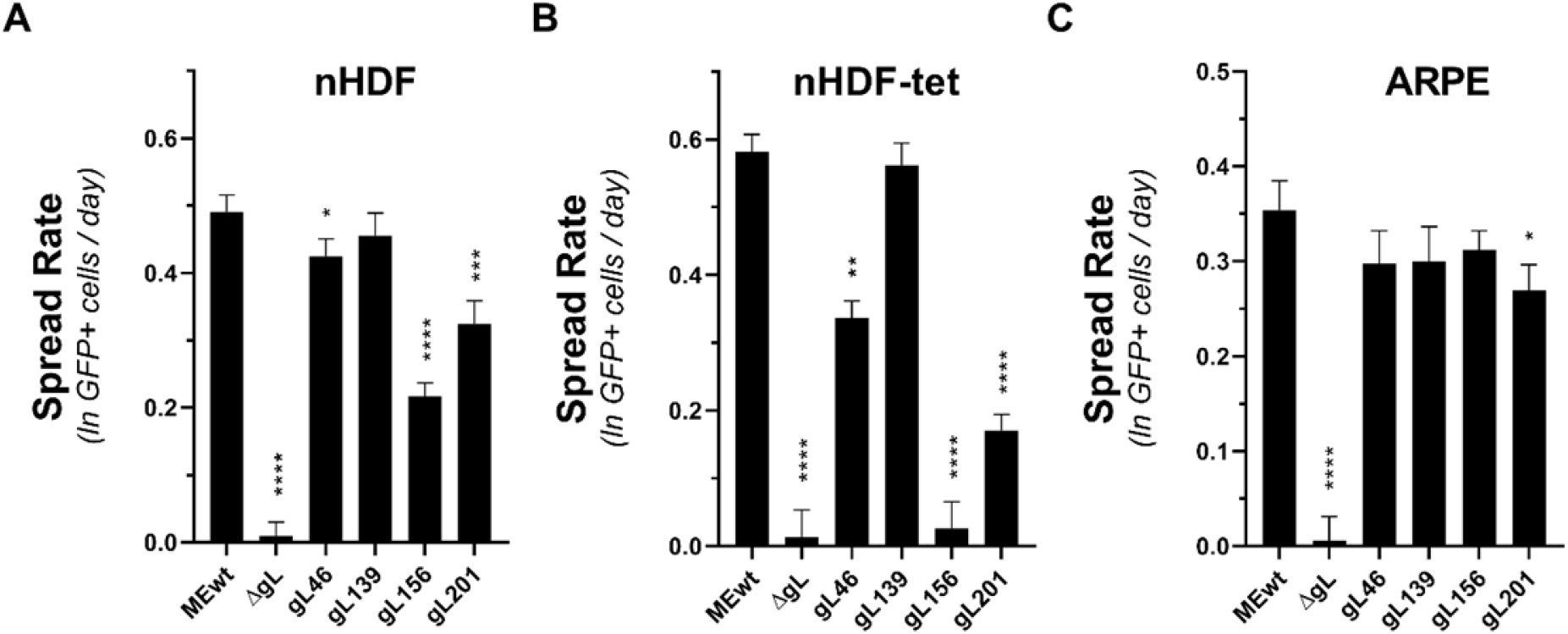
Effect of gL mutagenesis on spread of HCMV strain Merlin. nHDF (A), nHDF-tet (B), and ARPE19 (C) cells were infected at MOI 0.001 with HCMV strain ME or complemented ME gL mutants and spread efficiency was monitored over 12 days by flow cytometry. Average rates for three experiments are shown. All error bars represent standard deviation and P values were calculated using ANOVA with Dunnett’s multiple comparison test (*<0.05, **<0.01, ***<0.001, ****<0.0001).

## DISCUSSION

Since the initial characterizations of the UL128-131 proteins as important tropism factors (25– 28), much has been learned about the roles of gH/gL/gO and gH/gL/pUL128-131 including their partially overlapping requirements for entry and cell-cell spread of HCMV on different cell types, and the identification of multiple potential receptors for each, including PDGFRα, TGFβR3 for gH/gL/gO and integrins, NRP-2, and OR14I1 for gH/gL/pUL128-131 (16, 17, 20, 22, 24, 26–29, 47, 52). However, much still remains to be learned regarding the mechanisms by which the gH/gL complexes facilitate entry and spread and the specific roles of the receptor interactions. At a fundamental level, herpesvirus gH/gL complexes regulate membrane fusion through direct interactions with the fusion protein gB (15). For HSV and EBV, models suggest that receptor binding, either directly by gH/gL or through intermediary proteins like gD or gp42, induces conformation changes in gH/gL that promote interactions with gB, leading to fusion. Among the aforementioned HCMV receptors, PDGFRα is the only one for which there are direct data suggesting a role in the regulation of gB fusion activity during entry. Cell-cell fusion can be mediated by transient expression of gH/gL and gB, without gO or the UL128-131 proteins (40, 41). Vanarsdall et al. demonstrated that HCMV virion extracts contain stable gH/gL-gB complexes, and far less gB in stable complex with either gH/gL/gO or gH/gL/pUL128-131 (53). Subsequently, Wu et al., accounting for the results of Vanarsdall and the earlier evidence of interaction between PDGFRα and gB (54), suggested a stepwise model where the binding of gH/gL/gO to PDGFRα promotes the binding of gB to PDGFRα (55). In contrast, there have been no data indicating direct interactions between gH/gL/pUL128-131 and gB, but there is ample evidence that gH/gL/pUL128-131 can induce cell receptor-mediated signaling pathways that influence the nature of the entry pathway, whereas signaling through PDGFRα is not required for infection (20, 22, 24, 52, 55, 56). To further delineate the functions of gH/gL/gO and gH/gL/pUL128-131, we analyzed a library of gL CCTA mutants and found that several disproportionately affect the assembly and function of gH/gL/gO over gH/gL/pUL128-131.

In a previous study using Ad vector expression, we found that only 2 of the gL CCTA mutants (L139 and L244) were functional in a gH/gL-gB cell-cell fusion assay, but all were able to form gH/gL/pUL128-131 complexes that could induce receptor interference, suggesting a separation of the core fusion function of gH/gL from the receptor-binding capacity of gH/gL/pUL128-131 (41). To study how these mutations might distinguish gH/gL/gO and gH/gL/pUL128-131 during HCMV infection, we engineered the gL mutants into the genetic backgrounds of the HCMV BAC clones TB, TR, and ME, which differ in the expression of gH/gL/gO of gH/gL/pUL128-131, encode genetically distinct variants of gH and gO, and differ in their dependence on the gH/gL complexes for their mechanisms of spread (19, 34, 35, 57).

The most severe mutant spread phenotype was for L156, which nearly phenocopied a ΔgL mutant in gH/gL/gO-dependent spread conditions, i.e. spread in fibroblasts for TR, TB, and ME under gH/gL/pUL128-131 repression. By comparison, L156 had a far more moderate effect on ME when the robust expression of gH/gL/pUL128-131 was allowed to contribute, and no effect on spread in epithelial cells was observed. These data were consistent with our prior analyses indicating unimpaired gH/gL/pUL128-131 function (41). Mapping the L156 mutations on to the published structure models of gH/gL/gO and gH/gL/pUL128-131 (42, 58) suggests multiple stabilizing interactions with residues N179(gO) and N114(gL) for gH/gL/gO, and a single interaction with Q97 of pUL130 (Fig 8A). This would be consistent with the apparently more severe disruption in assembly of sgH/gL/gO complexes compared to sgH/gL/pUL128-131. Similarly, L201 disproportionately impaired gH/gL/gO-dependent spread over gH/gL/pUL128-131-dependent spread. While there was no obvious impact on the assembly of sgH/gL/gO for L201, there was substantially less gH/gL/gO in TB_L201 virions. However, two observations make it difficult to attribute all of the impaired infectivity and spread associated with the L201 mutation to the reduced amounts of gH/gL/gO in the virion. First, the L46 mutation also resulted in a dramatic reduction of gH/gL/gO in TB virions but had little or no effect on infectivity or spread. Second, L201 was the only mutant that formed sol. gH/gL/gO that was unable to block HCMV infection. Together, these observations indicate that L201 causes a functional disruption of gH/gL/gO beyond an assembly defect. The defect of L201 in the previous reported gH/gL-gB cell-cell fusion assay suggests impaired profusion interactions with gB, and inability of L201 containing sgH/gL/gO to block HCMV infection suggests important receptors beyond PDGFRα.

**Figure 8.**
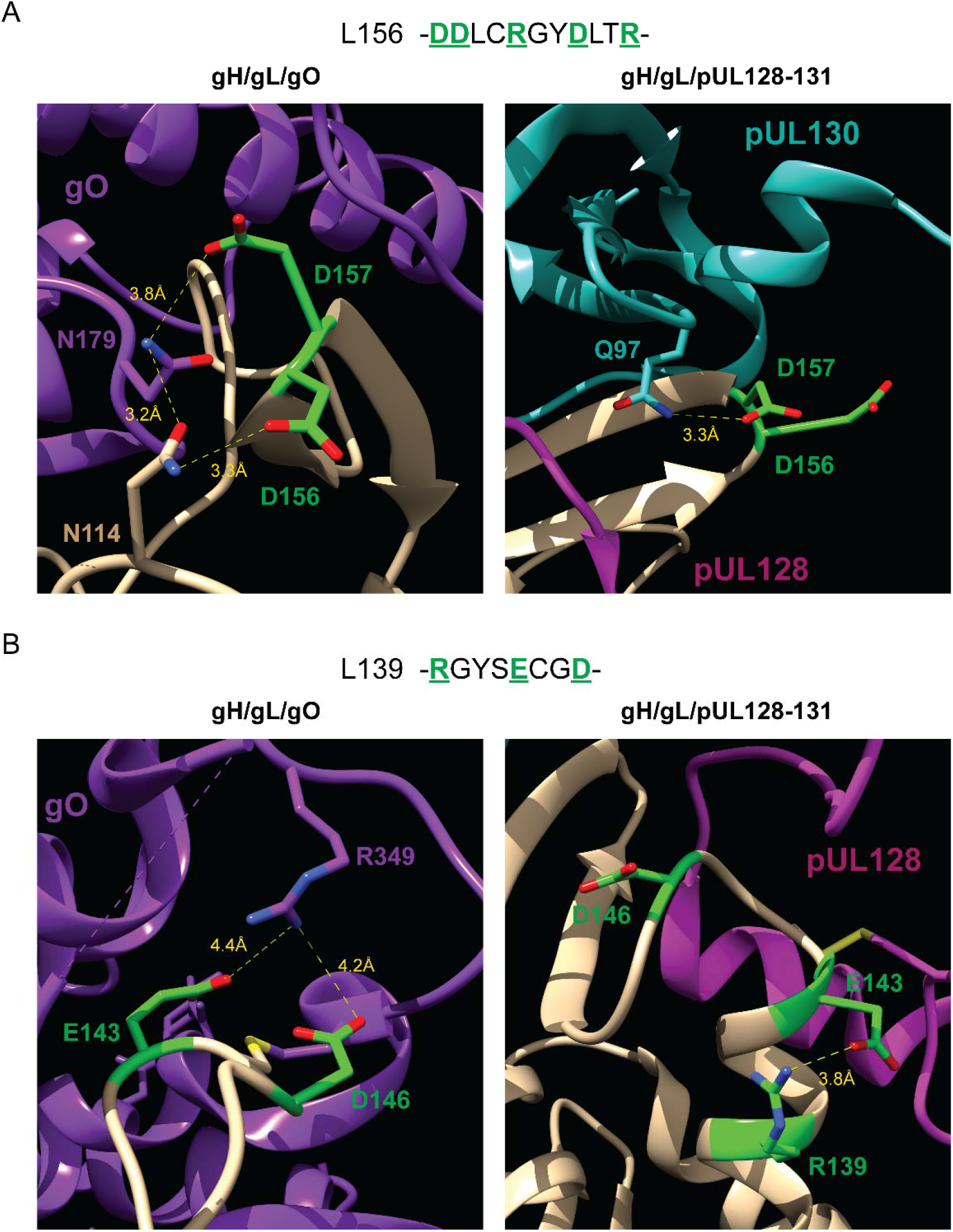
Structural differences between gH/gL/gO and gH/gL/pUL128-131 for CCTA regions L156 and L139. 3-D representations of the L156 (A) and L139 (B) regions containing CCTA mutations for gH/gL/gO (left panel) and gH/gL/pUL128-131 (right panel). Mutated gL residues of particular interest are colored green and electrostatic interactions are depicted with a yellow dashed line and corresponding distance (Å). All images generated in Chimera software using PDB codes 7LBE and 5VOB for gH/gL/gO and gH/gL/pUL128-131, respectively.

One striking result of these studies was that several of the gL mutants yielded viable HCMV, despite being inactive in the previous gH/gL-gB cell-cell fusion analyses (41). L46 stands out in this category, having caused only a minor reduction of spread for TR and virtually no reduction for TB. Like the other gL mutations, L46 did not significantly impact gH/gL/pUL128-131, inasmuch as there were no effects on spread of ME in fibroblasts with full gH/gL/pUL128-131 expression, or in epithelial cells. Thus, the minor effects observed for L46 were likely associated with gH/gL/gO function. The apparent discrepancy between the previous cell-cell fusion results and the viability of HCMV for gH/gL/gO-dependent spread might indicate that L46 disrupts some critical conformation of gH/gL that is restored by gO. Alternatively, this result might reflect fundamental differences between cell-cell fusion and virus-cell fusion. Moreover, it is notable that there were differences in the magnitude of the L46 effects between TR, TB, and ME under repressed gH/gL/pUL128-131, which point towards epistatic influences, potentially related to genetic variation in gH and gO among these strains.

L139 was the only mutant that enhanced any parameter measured. TR_L139 spread at a faster rate and displayed a markedly more diffuse focal pattern compared to the parental WT, suggesting an enhanced cell-free spread efficiency. However, TR_L139 virions were actually more than 10-fold less infectious. To our knowledge, this is the only example of such a discrepancy between cell-free spread efficiency and measured infectivity. Under the general herpesvirus model that activation of gB by gH/gL complexes involves structural rearrangements in gH/gL, it might be that the L139 mutation places gH/gL on a “hair trigger”, more prone to spontaneous (i.e., untriggered) conversion. If so, this might make the infectivity of TR_L139 virions more labile during harvesting and storing prior to the infectivity assays but may well result in enhancement of extracellular infectivity during the spread assays, which do not involve storage of the progeny virus. Although the previous gH/gL-gB cell-cell fusion assay was reported as a binary readout “fusion (+) or (-)”, L139 visually appeared to be “hyperactive”, and this would be consistent with the above idea of a “hair-trigger” ((41) and J.M. Lanchy; unpublished observations). Finally, as with L46, the effect of L139 on infectivity was not equally manifest in the different genetic backgrounds and had no effects on spread mediated by gH/gL/pUL128-131. The L139 cluster is noteworthy because it contains Cys144, the residue that makes a critical disulfide bond with pUL128. Mapping the L139 mutations to the published structures of gH/gL/gO and gH/gL/pUL128-131 (42, 58) offers potential structural explanations for disproportionate effects on gH/gL/gO and for the observed epistasis (Fig 8B). In gH/gL/pUL128-131 the L139 region forms a helix stabilized by the electrostatic interactions between R139 and E143, and likely provides stability for the disulfide interaction with pUL128. In contrast, the L139 region in gH/gL/gO is an unstructured loop with residues E143 and D146 make electrostatic interactions with R394 of gO. The loss of the interactions with gO could result in instability and explain the hyperactivity of L139 mutants. The unstructured nature of the L139 region in gH/gL/gO may be more prone to epistatic effects of sequence variation in gH and gO that lead to subtle conformational changes depending on the genetic background of HCMV.

The notion that gH/gL/pUL128-131 functions by promoting gB fusion activity is based on observations of efficient cell-to-cell spread by gO null HCMV. Both TB and TR gO null mutants were impaired for spread in fibroblasts, but spread as well or slightly better than their respective parental strains in endothelial or epithelial cells (16, 47, 55), whereas gO null ME spread comparably to the parental on both fibroblasts and endothelial cells (29). Complementary observations reported by Wu et al. showed that HCMV with intact gH/gL/pUL128-131 spread on PDGFRα-knockout fibroblasts, whereas HCMV lacking gH/gL/pUL128-131 could not (30). These observations indicate an important role for gH/gL/pUL128-131 in the spread route but offer little suggestion of the mechanism(s) involved. Moreover, synthesizing the results of these studies using distinct HCMV BAC clones into unified conclusion on the roles of gH/gL complexes in spread may be problematic. Our recent reports demonstrated that individual BAC clones can be highly specialized to one mode of spread, and suggested mechanistic differences between direct cell-to-cell spread and spread via extracellular virus (34, 35). All modes of HCMV spread likely require gB (59) and some kind of gH/gL complex, inasmuch as all gL null HCMV reported herein were unable to spread. However, ME is capable of one or more mechanisms that promote direct cell-to-cell spread that is far more efficient than that of TB, despite the low infectivity of the intracellular ME progeny virus (34). Moreover, some changes to the gH-gO allelic paring had effects on one mode of spread but not the other (35). Thus, the relative contributions of gH/gL complexes in direct cell-to-cell spread might differ significantly between strains, and may also differ between cell-to-cell spread and entry by extracellular virus.

Recent structures of gH/gL/gO and gH/gL/pUL128-131 indicate strong conservation of gH/gL structure underlying both complexes (42, 58). However, the results herein suggest that the functional regions of gL are different between the complexes. Collectively, our analyses support the model that gH/gL/gO can provide an activation signal for gB inasmuch as the mutations affected both gH/gL-gB cell-cell fusion (41) and gH/gL/gO-dependent spread of HCMV. Conversely, since the same gL mutations had little or no effect on gH/gL/pUL128-131-mediated receptor interference (41) or HCMV spread, the model that gH/gL/pUL128-131 provides a gB-activation function would seem to require different surfaces of gH/gL, implying a distinct mechanism of gH/gL/pUL128-131-triggered gB fusion.

## MATERIAL AND METHODS

### Cell lines

Primary neonatal human dermal fibroblasts (nHDF; Thermo Fisher Scientific), gL-nHDFs (nHDFs transduced with lentiviral vectors encoding codon-optimized UL115 of HCMV strain TR, selected with puromycin resistance), nHDF-tet (nHDFs transduced with retroviral vectors encoding the tetracycline repressor protein), gL-nHDF-tet, and MRC5 fibroblasts (American Type Culture Collection; CCL-171) were grown in Dulbecco’s modified Eagle’s medium (DMEM)(Sigma) supplemented with 5% heat-inactivated fetal bovine serum (FBS) (R&D Systems, Minneapolis, MN, USA) and 5% Fetalgro® (Rocky Mountain Biologics, Missoula, MT, USA), with penicillin-streptomycin, gentamicin, and amphotericin B. Retinal pigment epithelial cells (ARPE19)(American Type Culture Collection) were grown in a mixture of 1:1 DMEM and Ham’s F12 medium (DMEM:F12)(Sigma) supplemented with 10% FBS, with penicillin-streptomycin, and amphotericin B.

### Lentiviral and adenovirus vectors

The codon-optimized UL115 gene from HCMV strain TR (NCBI ref KF021605) was used to replace the eGFP ORF in the pLJM1-EGFP lentiviral transfer vector plasmid. The pLJM1-EGFP plasmid was a gift from David Sabatini (Addgene plasmid # 19319) (60). The gL-containing vector plasmid was transformed in 293T cells together with three lentiviral helper plasmids. The pMDLg/pRRE, pRSV-Rev, and pMD2.G helper plasmids were a gift from Didier Trono (Addgene plasmids # 12251, # 12253, # 12259, respectively) (61). Two days after transformation, the lentiviral particles in the supernatant were purified from cell debris thru syringe filtration and centrifugation. After titration, the particles were used to transduce either low passage nHDF or MRC-5 cells. After a week of puromycin selection, cells were tested for gL expression and aliquots were stored in liquid nitrogen until further usage.

Replication-defective (E1-negative) adenovirus (Ad) vectors that express HCMV TR sgH-6His, gO, UL128, UL130, UL131, and gL (wild type or bearing CCTA substitutions) were made as previously described (41). Briefly, Ad vector stocks were generated by infecting 293IQ cells at 0.1 PFU/cell for 6 to 10 days. The cells were pelleted by centrifugation, resuspended in DMEM containing 2% FBS, sonicated to release cell-associated virus, and cleared the cellular debris. Titers were determined by plaque assay on 293IQ cells. Multiplicities of infection (MOIs) for Ad vectors were determined empirically for each experiment and ranged from 3 to 30 PFU per cell. Because protein expression can vary between stocks of Ad vectors, experiments were performed with Ad vectors at different MOIs to account for the possible effects of under- or overexpression of proteins.

### HCMV

All human cytomegalovirus (HCMV) strains were derived from bacterial artificial chromosome (BAC) clones. BAC clone TR was provided by Jay Nelson (Oregon Health and Sciences University, Portland, OR, USA)(62). BAC clone TB40/e (BAC4) was provided by Christian Sinzger (University of Ulm, Germany)(12). BAC clone Merlin (pAL1393), which contains tetracycline operator sequences within the transcriptional promotor of UL130 and UL131, was provided by Richard Stanton (Cardiff University, United Kingdom)(33). UL115 mutants were generated by en passant recombineering (63) and modified to express GFP, as previously described (34). Infectious HCMV was recovered by electroporation into HFFFs (normal or gL-expressing), as previously described (16). For infectious unit determination, viruses were serial diluted, and infectivity was determined on fibroblasts using flow cytometry at 48h post infection. Multiplicities of infection were determined as IUs/cell.

### Antibodies

Monoclonal antibodies specific to HCMV major capsid protein (MCP) and gH (AP86) were provided by Bill Britt (University of Alabama, Birmingham, AL, USA) (64, 65). Rabbit polyclonal sera against HCMV gL, UL130, and UL131 were provided by David Johnson (Oregon Health Sciences University, Portland, OR, USA)(45). Monoclonal antibody specific to pUL128 (4B10) was provided by Tom Shenk (Princeton University, Princeton, NJ, USA)(46). Polyclonal antisera raised against peptides corresponding to residues 250 to 269 of TBgO was purchased from GenScript (Piscataway, NJ, USA.), as previously described (32). Anti-6His antibodies from mice (MA1-21315) were purchased from ThermoScientific.

### Expression of soluble gH/gL complexes

Soluble gH/gL complexes were expressed as previously described (43). Briefly, APRE19 cells were infected with replication-defective adenoviral vectors expressing soluble gH-6His (sgH), gL (mutant or WT), and gO or pUL128-131 for 24 hours. Inoculums were replaced with DMEM/F12 medium containing 2% FBS and collected after 7 days. Complexes were filtered with 0.22um SteriCup filter (Millipore), then enriched with Ni-NTA agarose resin (Thermo) and eluted with gel loading buffer absent of DTT. Total protein content was qualitatively assessed by precipitation using 80% acetone followed by immunoblot.

### HCMV inhibition

The inhibitory capacity of the soluble gH/gL/gO complexes was tested as described previously (48). The supernatants containing the soluble complexes were diluted with DMEM:F12 supplemented with 2% FBS. These dilutions, as well as the cells were precooled on ice for 15 min before the medium was removed from the cells and replaced by the dilutions of soluble gH/gL/gO, gH/gL, or medium as an untreated control. Incubation was performed at 4°C on ice to avoid receptor endocytosis as a mechanism of inhibition. The inoculum was aspirated and replaced by precooled TB40-BAC4 diluted to result in 50% infection. Binding of the virus to cells was performed for 1h at 4°C on ice before the cells were shifted 37°C to allow entry. After 4h at 37°C, the virus inoculum was removed and replaced by maintenance medium. After two days the cells were detached and fixed, and GFP+ cells were determined by flow cytometry.

### ELISA

Ni-NTA HisSorb plates (Qiagen) were incubated with soluble gH/gL/gO complexes tagged with 6-His overnight at 4°C. Medium only and supernatants from cells infected with Adenoviruses expressing only gH and gO (which do not secrete either of those proteins without gL) were used as negative controls. Unbound protein was removed by 4 rounds of washing with PBS + 0.05% Tween-20. Purified PDGFRα-Fc, generously provided by S. Feldmann (66), was diluted in PBS + 0.2% BSA and incubated on the plates for 2h at room temperature. Unbound soluble receptor was removed by washing 4 times with PBS + 0.05% Tween-20. The secondary goat anti-human HRP antibody (Invitrogen A18817) was diluted to 100 ng/ml in PBS 0.2% BSA and incubated on the plates for 45min at room temperature. After washing twice with PBS 0.05% Tween-20 and twice with PBS, TMB substrate was added and incubated for 15min before the reaction was stopped by addition of 2 mol/l sulfuric acid. The absorbance was measured at 450 nm and corrected for background signals as determined with the conditioned control.

### Flow Cytometry

GFP-expressing HCMV-infected cells were washed twice with PBS and lifted with trypsin. Trypsin was quenched with DMEM containing 10% FBS and cells were spun at 500 x g for 5 minutes at RT. Cells were fixed in PBS containing 2% paraformaldehyde for 10 min at RT, then washed and resuspended in PBS. Samples were analyzed with an AttuneNxT flow cytometer, as previously described (34). HCMV-infected cells were gated first by FSC-A and SSC-A, then for single cells using FSC-W and FSC-H, and GFP+ cells were measured with the BL1-A laser (488nm).

### qPCR

Viral genomes were determined as previously described (35). Briefly, supernatants containing cell-free HCMV were treated with DNase I, then viral genomic DNA was extracted using the PureLink viral RNA/DNA minikit (Thermo). Primers specific to sequences with UL83 were used with the MyIQ real-time PCR detection system (Bio-Rad).

### Immunoblot analysis

Infected cells, cell-free virions, and soluble gH/gL complexes were solubilized in a buffer containing 20 mM Tris-buffered saline (TBS) (pH 7.2) and 2% SDS. For reducing conditions, DTT was added just prior to analysis. Protein samples were separated by SDS-polyacrylamide gel electrophoreses (SDS-PAGE) and electrophoretically transferred to polyvinylidene difluoride (PVDF) membranes in a buffer containing 10 mM NaHCO3 and 3mM Na2CO3 (pH 9.9) and 10% methanol. Transferred proteins were first probed with MAbs or rabbit serum, then anti-mouse or anti-rabbit secondary antibodies conjugated to horseradish peroxidase (Sigma), and Pierce ECL Western blotting substrate (Sigma). For reprobing, antibodies were stripped from membranes using 25mM glycine with 1% SDS, pH2, washed, and then re-incubated with secondary-HRP and imaged to verify no residual chemiluminescent signal. Chemiluminescence was detected using a Bio-Rad ChemiDoc MP imaging system.

### Statistical analysis

Unless otherwise stated, all experiments were performed a minimum of three times. All curve fitting and statistical analysis was done using GraphPad Prism 9 software. Experiments comparing multiple mutants were analyzed by ANOVA with Dunnett’s multiple comparisons test (95% CI). Standard two-tailed *t* tests were used for direct comparisons. Error bars represent standard deviation between experiments, and *p*-values are represented as follows: *, *p* < 0.05; **, *p* < 0.01; ***, *p* < 0.001; ****, *p* < 0.0001.

## ACKNOWLEDGEMENTS

We are grateful to Bill Britt, Jay Nelson, Christian Sinzger, Richard Stanton, and David Johnson for generously supplying HCMV BAC clones, antibodies, soluble PDGFRα-Fc, and cell lines, as indicated in Materials and Methods. Additionally, we are grateful to the Center for Biomolecular Structure and Dynamics (CBSD), University of Montana, Missoula, MT, for purification of monoclonal antibodies and ELISA instrumentation, as well as the Flow Cytometry Core of the Center for Environmental Health Sciences (CEHS), University of Montana, Missoula, MT, for guidance on experimental design, acquisition, and analysis of the flow cytometry-based approaches used for this study.

This work was supported by a grant from the National Institutes of Health (NIH) to B.J.R (R01AI097274), a fellowship from the American Heart Association (AHA) to E.P.S. (17POST33350043), a fellowship from the German Research Foundation (DFG) to C.S. (STE 2835/1-1), a NIH CoBRE award to the Center for Biomolecular Structure and Dynamics at University of Montana (PG20GM103546), and by an Institutional Development Award (IDeA) from the National Institute of General Medical Sciences of the NIH to the Center for Environmental Health Sciences at University of Montana (P30GM103338).

Experiments were designed by E.P.S., C.S., L.Z.-D., B.J.R., and J.-M.L. and performed by E.P.S., C.S., and Q.Y., and the manuscript was prepared by B.J.R., E.P.S., C.S., and J.-M.L.

